# Labeling Natural Killer cells with superparamagnetic iron oxide nanoparticles for detection by preclinical and clinical-scale magnetic particle imaging

**DOI:** 10.1101/2024.03.08.583881

**Authors:** Olivia C. Sehl, Yanwen Yang, Ariana R Anjier, Dmitry Nevozhay, Donghang Cheng, Kelvin Guo, Benjamin Fellows, A. Rahman Mohtasebzadeh, Erica E. Mason, Toby Sanders, Petrina Kim, David Trease, Dimpy Koul, Patrick W. Goodwill, Konstantin Sokolov, Max Wintermark, Nancy Gordon, Joan M. Greve, Vidya Gopalakrishnan

**Author notes:** Correspondence: Vidya Gopalakrishnan, Ph.D. Department of Pediatrics 1515 Holcombe Blvd, Houston, Texas 77030, USA Olivia C. Sehl, Ph.D. Magnetic Insight Inc. 2020 N Loop Rd, Alameda CA 94502, USA.

## Abstract

**Introduction:** Clinical adoption of NK cell immunotherapy is underway for medulloblastoma and osteosarcoma, however there is currently little feedback on cell fate after administration. We propose magnetic particle imaging (MPI) for the detection, localization, and quantification of VivoTrax-labeled NK cells.

**Methods:** Human-derived NK-92 cells were labeled by co-incubation with VivoTrax for 24 hours then the excess nanoparticles were washed with centrifugation. Cytolytic activity of labeled vs. unlabeled NK-92 cells was assessed after 4 hours of co-incubation with medulloblastoma cells (DAOY) or osteosarcoma cells (LM7 or OS17) using bioluminescent or GFP counts. Labeled NK-92 cells at two different doses (0.5 or 1 x 10^6^) were administered to excised mouse brains (cerebellum), tibias, and lungs then imaged by 3D preclinical MPI (MOMENTUM imager) and localized relative to fiducial markers. NK-92 cells were imaged by clinical-scale MPI under development at Magnetic Insight Inc.

**Results:** NK-92 cells were labeled with an average of 3.17 pg Fe/cell with no measured effects on cell viability or cytolytic activity against 3 tumor cell lines. MPI signal was directly quantitative with the number of VivoTrax-labeled NK-92 cells, with preclinical limit of detection of 3.1 x 10^4^ cells on MOMENTUM imager. Labeled NK-92 cells could be accurately localized in mouse brains, tibias, and lungs within < 1 mm of stereotactic injection coordinates with preclinical scanner. Feasibility for detection of a clinically relevant dose of 4 x 10^7^ labeled NK-92 cells was demonstrated on clinical-scale MPI.

**Conclusion:** MPI can provide sensitive, quantitative, and accurate spatial information on NK cell delivery, showing its potential to resolve a significant unmet clinical need to track NK cell treatments in patients.

## Introduction

Medulloblastoma (MB) and osteosarcoma (OS) are the most common malignant brain tumors and bone tumors in children and young adults.^1–3^ The prognosis for patients with recurrent/refractory and metastatic medulloblastoma continues to be poor and efforts over the past 30 years to develop more effective treatments have met with limited success. Despite aggressive chemotherapy and successful control of primary OS tumors, the 5-year survival for patients with resectable, localized disease remains at 60-70% and for patients with metastatic disease either at diagnosis or relapsed is less than 30%.^2^ Pulmonary metastasis remains the main cause of death in these patients with OS^4^. Thus, new therapeutic strategies are needed to improve outcomes for patients with recurrent, metastatic, and resistant disease in these patients.

Immunotherapy represents an important therapeutic option for recurrent, metastatic, and potentially unresectable solid tumors. NK cells play a crucial role in the innate immune response to viruses and tumors. Unlike T cells, NK cell cytolytic activity is not reliant on antigenic stimulation, which makes it particularly relevant to pediatric cancers where the antigenic landscape is not well defined^5–9^. NK cells induce tumor lysis through a balance of signaling activity between activating ligands and inhibitory receptors and ligands^10^. In addition, NK cells are not prone to causing cytokine storm, neurotoxicity, graft versus host disease, or indefinite expansion *in situ*.^11^ NK cell cytotoxicity is mostly mediated by the secretion of cytokines and effector molecules such as interferon gamma (IFNγ), interleukin 2 (IL-2), interleukin 12 (IL-12), interleukin 15 (IL-15), tumor necrosis factor alpha (TNF α) and interleukin 21 (IL-21).^12^ NK cells have demonstrated potent treatment responses in several malignancies, including sarcomas, carcinomas, lymphomas, and leukemias.^13–18^ This means that allogeneic adoptive cell therapy (ACT) involving NK cells could be used in many patients without requiring modification of a patient’s own cells as is currently necessary with CAR T cells (autologous ACT). Researchers at MD Anderson Cancer Center have successfully developed a platform technology for the clinical translation for *ex vivo* expansion of NK cells in sufficient therapeutic quantities from cord blood and other donors.^19,20^ Unfortunately, other barriers exist including lack of tools to follow NK cells trafficking, growth and proliferation within the tumor microenvironment (TME) and immune evasion, which could potentially prevent them from executing their biological function. This highlights the need for ACT-compatible non-invasive imaging tools to longitudinally follow cell therapy fate, especially in the context of solid tumors, to assess correlation with efficacy.

Our current approach is to monitor for NK cell treatment progression using anatomic imaging modalities such as magnetic resonance imaging (MRI) and computed tomography (CT)^21^. Because immunotherapy can frequently cause a responder’s tumor to grow in size before shrinking, post-treatment imaging results remain difficult to interpret^22^. For brain tumors, cerebrospinal fluid/blood draw is used to monitor the loco- regional presence of NK cells in tumors and to correlate with tumor size and patient survival. For OS, changes in the immune milieu in peripheral blood is used to correlate with tumor size and patient survival. However, this in not reflective of actual changes in the TME and in the absence of information on NK cell localization within the tumor, the lagging indicator of tumor size are indirect measures of NK cell activity at best, and difficult to interpret.

To date, there are no reports of cellular tracking of NK cells in clinical trials. Other cell types such as dendritic cells, stem cells, and pancreatic islet cells have been tracked in humans using iron-based magnetic resonance imaging (MRI)^23^, with the most recent study in 2015.^24^ Unfortunately, the “negative” signal contrast created by superparamagnetic iron oxide nanoparticles in MRI is non-specific and non-quantitative, as the signal void can be confused with voids and other sources of negative contrast. To overcome this, ^19^F agents were developed that create a hotspot signal that can be overlaid with a ^1^H anatomical image.^25,26^ Despite some encouraging results from our group and others, the low sensitivity is limiting pre-clinical and clinical use.^27–29^ Nuclear imaging techniques (PET and SPECT) produce excellent image contrast that is quantitative but can be detrimental to cell viability.^30^ Magnetic particle imaging (MPI) is an emerging modality for *in vivo* tracking of fate, biodistribution, quantification, and persistence of cellular therapeutics. To enable cell detection by MPI, cells are pre-labeled in culture with iron oxide nanoparticles. Most MPI cell tracking studies have been performed with adherent cell types, such as stem cells^31,32^, dendritic cells^33,34^, cancer cells^35^, and macrophages^36^. There is a growing desire to track T cells and NK cells, which unlike stem, dendritic and cancer cells, are non-adherent (suspension) cells^37,38^. These pose additional challenges such as: first, suspension cells tend to be less phagocytic to iron oxide nanoparticles leading to lower MPI sensitivity, and second, it is more challenging to eliminate free iron from suspension cells post-labeling. To address these challenges, pre-clinical development of a protocol to label NK cells with iron oxide nanoparticles without affecting cell viability or function is needed. There are several human-scale MPI scanners under development worldwide^39–42^, including a general-purpose human MPI scanner at Magnetic Insight Inc.^43^

In this work we aim to demonstrate a novel magnetic particle technology for the tracking of therapeutic NK cells post administration. We label human-derived NK cells with VivoTrax (Magnetic Insight Inc., Alameda, CA) for detection by preclinical and clinical-scale MPI. We directly compare NK cell labeling and MPI detection with two other cell types: T cells (suspended cell line) and phagocytic macrophages (adherent cell line). We assess viability of 3 cell types and cytolysis activity of NK cells after labeling. Further, we demonstrate precise 3D localization and quantification of labeled NK cells administered to mouse brains, tibias, and lungs using preclinical MPI, relevant to detection of NK cells for medulloblastoma, osteosarcoma, and lung metastases, respectively. Finally, we demonstrate the feasibility for detection of clinically relevant doses of labeled NK cells by human-scale MPI.

## Methods

### Cell culture

NK-92 cells (CRL-2407, ATCC) were cultured in Myelocult^TM^ H5100 media (StemCell Technologies) supplemented with 100 units/mL recombinant IL-2 IS (130- 097-744, Miltenyi) and 11% horse serum (26050088, Gibco). Jurkat T cells (TIB-152, ATCC) were cultured in RPMI-1640 media (30-2001, ATCC). RAW 264.7 macrophages (TIB-71, ATCC), DAOY medulloblastoma cells (HTB-186, ATCC), and osteosarcoma cell lines (LM7, OS-17) were cultured in DMEM media (30-2002, ATCC). Cells were grown at 37°C and 5% CO_2_.

### Cell labeling procedure

NK-92 cells were labeled by co-incubation of 0.6 x 10^6^ NK-92 cells in 500 mL serum-free media containing 50 µg or 200 µg Fe/mL VivoTrax (Magnetic Insight Inc., Alameda, CA) magnetic particles for 4 hours. After 4 hours, an equal volume of serum-containing medium was added to cell culture, for the remaining 16-hour incubation with VivoTrax. Cell-free (sedimentation) controls contained 50 µg, or 200 µg Fe/mL VivoTrax in the same volume of media without NK-92 cells.

For comparison of cell labeling across multiple cell types, Jurkat T cells were labeled with VivoTrax with the same protocol using 200 µg Fe/mL VivoTrax. Likewise, RAW 264.7 macrophages were labeled with 200 µg Fe/mL VivoTrax and were seeded the day prior to allow adherence of cells to the flask for labeling.

At the end of the 24-hour co-incubation period, cells were washed to remove excess iron oxide nanoparticles. NK-92 and Jurkat T cell suspensions were transferred to 15 mL Falcon tubes. To obtain cell suspension of adherent RAW 264.7 macrophages, cells were detached using trypsin (0.25% trypsin-EDTA, 25200056, Gibco). Cells were suspended in 10 mL serum-containing media, mixed gently, and centrifuged at 300 g for 8 minutes. This washing procedure was repeated a total of 3 times to remove extracellular iron from labeled cells. Sedimentation (no cell) controls were washed by the same protocol. Cell counting were performed with Cellometer (Nexcelom Bioscience) or Vi-cell (Beckman) counters then cells were used for cytology and cell staining, prepared as cell pellets for determination of iron labeling and MPI limit of detection, or used for cell functional assays.

### Assessment of cell labeling

For qualitative cell labeling assessment using histology, VivoTrax-labeled NK-92 cells were centrifuged onto Cytospin glass slides at a density of 1 x 10^5^ cells (Cytospin 4 centrifuge, Thermo Scientific). Attached cells were then fixed by methanol/acetic acid solution (3:1) stained with Perl’s Prussian blue with nuclear fast red as counterstain (Iron stain kit, ab150674, abcam). Slides were dehydrated (incubation with 75, 95, and 100% ethanol) and incubated with xylene before coverslipping. Cytospin slides were imaged using an Olympus IX81 microscope with 20x objective lens. The percentage of labeled cells was calculated as (number of cells with iron staining) / (total number of cells) * 100% and was measured from 7 FOVs per slide.

For quantitative cell labeling assessment using MPI, NK-92 cell pellets containing 1 x 10^6^ cells were prepared for each VivoTrax concentration (50 vs. 200 µg Fe/mL). The equivalent volume of suspension from sedimentation controls were also prepared. NK- 92 cell pellets were imaged on a MOMENTUM^TM^ preclinical scanner (Magnetic Insight) using the following parameters: 2D field of view (FOV) = 6 x 6 cm, transmit axes = x and z (multichannel imaging), gradient strength = 3.055 T/m, RF amplitude = 20 mT (x-axis) and 23 mT (z-axis), excitation frequency = 45 kHz, averages = 1. To determine cellular detection limit for preclinical MPI, a dilution series of VivoTrax-labeled NK-92 cells, Jurkat T cells, and RAW 264.7 macrophages were prepared by serial 1:1 dilutions from 1 x 10^6^ cells. These cell pellets were imaged on preclinical MOMENTUM imager using the same parameters. Analysis and quantification of MPI signal for cell pellets and sedimentation controls is described below.

For quantitative cell labeling assessment using colorimetric iron assay kit, 2 x 10^6^ labeled NK-92 cells, Jurkat T cells, and RAW 264.7 macrophages were digested with 5 mL concentrated (67-70%) nitric acid and heated to 120 °C for complete evaporation of acid. The remaining iron nitrate salt was allowed to cool then quantified using a colorimetric iron assay kit with 3 technical replicates for each cell type (MAK025, Sigma). The total iron from 3 technical replicates was averaged then divided by the number of cells to obtain calculated pg Fe/cell.

### Measurement of NK cell activity

NK-92 cell viability was determined using trypan blue exclusion assay and Vi-cell counter (Beckman). For NK-92 cytotoxicity assay with DAOY medulloblastoma cells, DAOY cells were engineered to stably express firefly luciferase (ffluc) then were seeded in a 96 well plate (10,000 cells /well) overnight. Unlabeled or VivoTrax-labeled NK-92 cells (at concentration 200 µg Fe/mL) were added at various effector-to-target (E/T) ratios to interact at 37°C for 4 h. A final concentration of 0.15 mg/ml D-luciferin (PerkinElmer; #122799) was added to each well for analysis of surviving cell. The plate was read by a CLARIOstar Plus multi-mode plate reader (Cary, NC, USA) for luminescence counts per second (relative light units, RLU).

For the NK-92 cell cytotoxicity assay with osteosarcoma cells, LM7-GFP and OS17-GFP cells were seeded in a 96 well plate (10,000 cells/well). Unlabeled or VivoTrax-labeled NK-92 cells (at concentration 200ug Fe/mL) were added at various effector-to-target ratios to interact at 37°C for 4 h. The green fluorescence of LM7-GFP and OS17-GFP cells was measured using an IncuCyte® S3 Live-Cell Analysis Instrument. Green fluorescent area (in μm^2^) was compared across the various effector-to-target ratios. These measurements were also compared with spontaneous death (media) and maximum death (2% Triton) controls to calculate Percent Specific Lysis using the following formula: [(spontaneous death GFP area – test GFP area)/(spontaneous death GFP area – maximum death GFP area)]*100.

Proliferation of Jurkat T cells, and raw264.7 macrophages was assessed using quantitative colorimetric MTS assay (ab197010, Abcam), after incubation with a range of VivoTrax concentrations (0-400 µg Fe/mL) for 24 hours. This assay was conducted in accordance with International Organization for Standardization (ISO) documentation 19007.^44^

### Implantation of NK cells in mouse organs

To assess feasibility for NK-92 cell detection with MPI for medulloblastoma and osteosarcoma, NK-92 cells were implanted to mouse cerebellum or tibias. C57BL/6 mice of approximately 6-8 months were euthanized and their brains and tibias were harvested prior to cell implantation. Further, to represent treatment for osteosarcoma lung metastasis, NK-92 cells were implanted into the lungs of euthanized mice, as described below. A summary of our NK-92 cell implantations to mouse organs is outlined in **Table 1** and described in detail below. Implanted NK-92 cells were unlabeled (control) or labeled using the protocol described above with a concentration of 200 µg Fe/mL. An equal number of male and female mice organs were used for this study.

**Table 1.**
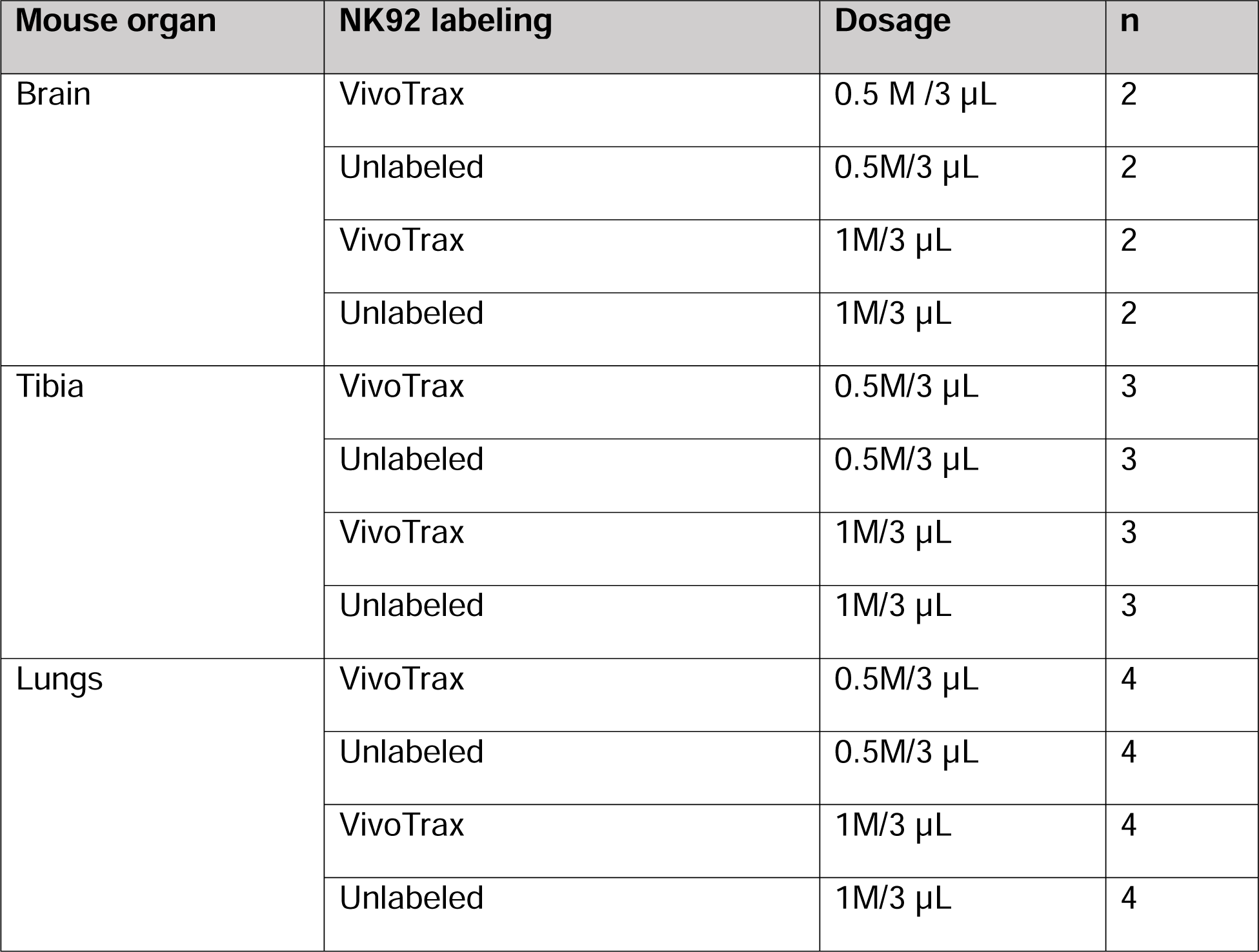
Summary of mouse organs from euthanized C57BL/6 mice and injection strategy. For each organ, the NK-92 cell labeling, cell dose (M = million), and sample size are tabulated.

For brains, we implanted 5 x 10^5^ NK-92 cells (n = 2 VivoTrax labeled; n = 2 unlabeled) or 1 x 10^6^ NK-92 cells (n = 2 VivoTrax labeled; n = 2 unlabeled) suspended in 3 µL saline. NK-92 cells were implanted in the mouse cerebellum with coordinates: 2 mm lateral (right) and 2 mm caudal of lambda using a stereotaxic device and a 10 μL Hamilton syringe. The needle was inserted at a depth of 2.5 mm, and the cell solution was injected at a rate of 1.0 µL/min.

For legs, NK-92 cells were injected through the proximal end of the tibia. Injections consisted of 5 x 10^5^ NK-92 cells (n = 3 VivoTrax labeled; n = 3 unlabeled) or 1 x 10^6^ NK-92 cells (n = 3 VivoTrax labeled; n = 3 unlabeled) suspended in 3 µL saline. The skin and hair were left intact on the removed legs during the injection. A 30-gauge needle was used to gently drill through the kneecap of the dissected leg and then the needle was pulled back slightly to create a small space for injected NK-92 cells into the tibia with the goal of injecting the cells into the middle of the tibia.

For lungs, VivoTrax-labeled NK-92 cells were injected into the dissected lungs on the left side from a euthanized mouse through the primary bronchus, with unlabelled NK-92 cells injected into the contralateral side of the lungs (right side). The heart was dissected along with the lungs to allow for orientation of the organs. To prevent the cells from flowing out of the lungs or into the other side of the lungs, a knot was tied around the left and right primary bronchi where they meet the trachea which allowed for separation of the two sides of the lungs for VivoTrax-labeled and unlabeled NK-92 cells to be compared. Differently colored thread was used to identify which side received unlabeled vs. VivoTrax- labeled NK-92 cells. This was repeated for 0.5 x 10^6^ cells (n = 4 lungs) or 1.0 x 10^6^ cells (n = 4 lungs).

Immediately after NK-92 cell injections to brains, lungs, and legs, organs were placed into individual tubes and flask frozen in liquid nitrogen. Frozen organs were shipped to Magnetic Insight for MPI. In a separate experiment, we studied MPI signal stability with a freeze-thaw cycle for frozen VivoTrax and VivoTrax-labeled cells (**Supplementary** Figure 1).

### MPI of mouse organs

MPI was performed for each mouse organ on a MOMENTUM MPI scanner to assess cell location. At the beginning of the imaging session, a 2D image was acquired of the empty sample holder to assess background signal levels. Each mouse organ was imaged individually by placement in the center of the imaging holder. MPI projections were acquired in 2D with the following parameters: FOV = 12 x 6 cm, transmit axes = x and z (multichannel imaging), gradient strength = 5.7 T/m, RF amplitude = 20 mT (x-axis) and 23 mT (z-axis), averages = 1. 3D imaging was performed with the following parameters: FOV = 12 x 6 x 6 cm, 35 projections, and reconstruction using Filtered Back Projection. During 3D imaging, three fiducials with 10% VivoTrax (in 1 µL) were included to localize NK-92 cell signal in tissue.

### Imaging NK cells on clinical-scale MPI

For phantom testing at the human scale, a clinically relevant dose (40 x 10^6^) of NK-92 cells was labeled using the labeling protocol with 200 µg Fe/mL media and then pelleted in a 15 mL Falcon tube. These NK-92 cells were imaged using a clinical coil with inner diameter (ID) = 25 cm. The imaging pulse sequence scans a 20 cm (x) x 20 cm (y) x 3 cm (z) Field of View and has an image acquisition time of 5.5 minutes.

For sensitivity limit studies, a small detection coil (ID = 2 cm) was used. Cell pellets imaged with the coil included: 1 x 10^6^ VivoTrax-labeled NK-92 cells labeled with 50 µg Fe/mL (n = 3 pellets) or 200 µg Fe/mL (n = 3 pellets), and dilution series of T cells and macrophages. The imaging pulse sequence scans a 20 cm (x) x 20 cm (y) x 1 cm (z) Field of View and has an image acquisition time of 6.4 minutes.

The clinical-scale MPI device used in this experiment has a 60 cm magnet-free bore and produces images using a field free point (FFP) with a 0.28 x 0.28 x 0.55 T/m gradient strength.

### MPI analysis and quantification

We analysed 2D MPI images to quantify MPI signal from cell pellets (NK-92, T cells, and macrophages) and mouse organs (brains, legs, lungs). MPI dicom files were analyzed in MagImage software (Magnetic Insight, Inc.). For display, an MPI color bar was applied to images. 2D projection images were windowed/leveled to a uniform range for each organ to visualize differences in MPI signal due to varying number of injected cells. For MPI signal quantification, the standard deviation of background levels was determined (SD_bkg_) from image(s) of the empty sample holder. A threshold of 5 * SD_bkg_ was used to select and then sum signals from cells/organs imaged with 2D MPI.^45^ If there was no signal exceeding this threshold (e.g., control organs that received unlabeled cells), MPI signal was measured from a circular region of similar volume and localized over the region where the organ was positioned.

We analysed 3D MPI images to measure the location of cell implantation to mouse brains, legs, or lungs. 3D images were displayed as maximum intensity projections (MIP) along with individual slice views for each of three anatomical planes, using standard radiologic convention for orientation. The distances between maximum signals from fiducials and NK-92 cell signal were measured to calculate and verify the injection site. The MPI analyst was blinded to the stereotactic coordinates for NK-92 cell injection.

### Statistics

Unpaired t-tests were used for cytotoxicity assay to compare RLU from DAOY cells across NK-92 cell labeling (labeled vs. unlabeled) (n = 3). For each mouse organ, one-way analysis of variance (ANOVA) with Tukey’s multiple comparisons test was used to compare MPI signal measured from different cell numbers (unlabeled NK-92 cells vs. 0.5x10^6^ labeled NK-92 cells vs. 1x10^6^ labeled NK-92 cells). Statistical comparisons were considered significant if *p* < .05.

## Results

### *In vitro* analysis of NK-92 cell labeling

To qualitatively assess iron labeling of NK-92 cells, we performed Perl’s Prussian blue staining and verified the presence of iron within NK-92 cells at both labeling concentrations (50 and 200 µg Fe/mL) (**Fig 1.a**). Approximately 11% of NK-92 cells were Prussian blue positive at 50 µg Fe/mL and 26% of NK-92 cells were Prussian blue positive at 200 µg Fe/mL.

**Figure 1.**
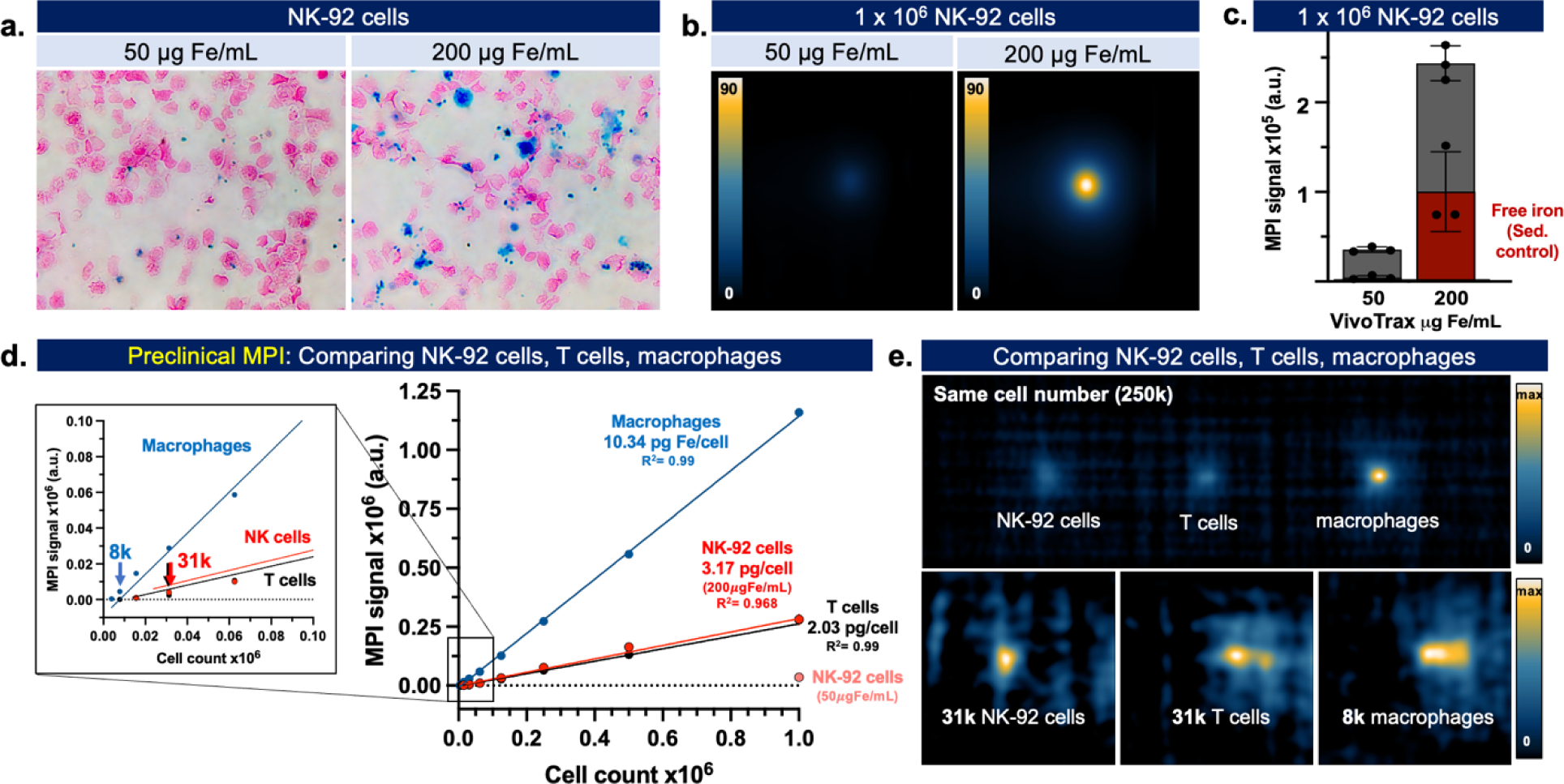
**a.** Perl’s Prussian blue (PPB) staining showed positive iron staining associated with VivoTrax-labeled NK-92 cells using both labeling concentrations. **B**. Comparing MPI signal from VivoTrax-labeled NK-92 cell pellets (1 x 10^6^ cells), labeled at 50 or 200 µg Fe/mL. **c**. MPI signal quantification shows higher signal from NK-92 cells labeled at 200 µg Fe/mL compared to 50 µg Fe/mL. However, there was also increases in MPI signal from sedimentation (sed) controls used to estimate free iron, as shown in red. **D.** A dilution series of VivoTrax-labeled NK-92 cells, T cells, and macrophages labeled using the same protocol imaged by MPI. The limit of detection was determined as 31x10^3^ (31k) for NK-92 cells and T cells and 8x10^3^ (8k) for macrophages. For each cell type, cell number was directly linear with MPI signal produced (R^2^ > 0.96). **e. [top]** MPI of 250k NK-92 cells, T cells, and macrophages for visual comparison. **[bottom]** MPI of the lowest detected number NK-92 cells (31k), T cells (31k), and macrophages (8k).

MPI of NK-92 cell pellets (1 x 10^6^ cells) is shown in **Fig. 1b**. On average, integrated MPI signal was 6.8x higher for NK-92 cells labeled at 200 µg Fe/mL compared to 50 µg Fe/mL (**Fig. 1c**). Sedimentation controls indicate approximately 10% of the MPI signal was associated with extracellular iron at the 50 µg Fe/mL condition. At 200 µg Fe/mL, the sedimentation control indicated that approximately 41% of MPI signal was associated with extracellular iron. As determined by the quantitative iron assay kit, average NK-92 cell labeling was 3.17 pg/cell. For comparison, macrophage labeling was 10.34 pg/cell and Jurkat T cell labeling was 2.03 pg/cell, obtained using the same labeling protocol.

The lowest number of NK-92 cells detected by preclinical MPI was 3.1 x 10^4^ cells. Comparatively, the fewest number of T cells detected was 3.1 x 10^4^ cells and the fewest number of macrophages detected was 8 x 10^3^ cells (**Fig. 1d**). MPI signal was directly linear with cell number for NK-92 cells, T cells, and macrophages (R^2^ > 0.96). The slope of the line of best fit for macrophages was 4x higher than NK-92 cells and 4.4x higher than T cells, reflecting cellular uptake by these cell types. Demonstration of preclinical MPI detection of the lowest cell numbers is shown in **Fig. 1e**.

### NK-92 cell activity

It is essential to assay cell viability and function to ensure that VivoTrax labeling does not hinder NK-92 cell activity. Analysis by trypan blue assay indicated NK-92 cell viability was 93.1% after 24h co-incubation with 50 µg Fe/mL VivoTrax and 91.2% at concentration 200 µg Fe/mL (average of n = 3 technical replicates). Assessment of cell function is demonstrated in **Figure 2**. Cytolysis assay shows there were no significant differences in unlabeled vs. VivoTrax-labeled NK-92 cytolysis against DAOY medulloblastoma cells, LM7 osteosarcoma cells, or OS17 osteosarcoma cells at various ratios (n.s., p > .05) (**Fig 2a**). MTS viability assay shows no significant differences in the proliferative capacity of Jurkat T cells, and mouse macrophages following 24-hour co-incubation with VivoTrax concentrations used for cell labeling (0-200 µg Fe/mL). A minimum of 80% proliferation is retained for cells incubated with VivoTrax compared to unlabeled cells.

**Figure 2:**
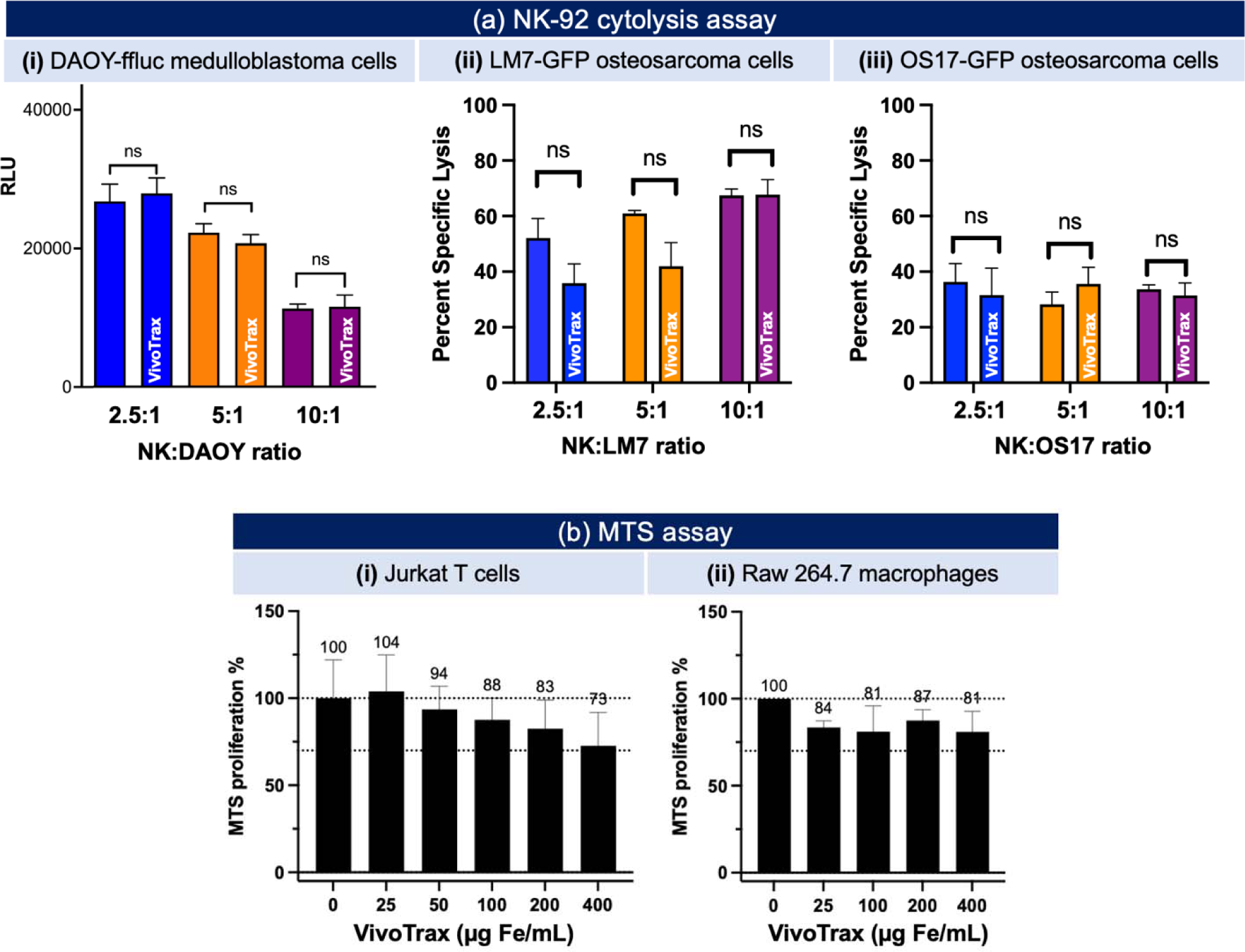
(a) NK-92 cytolysis assay with (i) DAOY-ffluc cells, (ii) LM7-GFP, and (iii) OS17-GFP cells. Unlabeled or VivoTrax-labeled NK-92 cells (effector) were added to target cells at effector:target ratios 2.5:1, 5:1, 10:1. Luciferase or GFP activity in surviving target cells was determined after 4 h of incubation. Data presented as mean with standard deviation. (c) MTS assay for Jurkat T cells and raw264.7 macrophages after 24 h co-incubation with VivoTrax particles at 0 – 400 µg Fe/mL. Data presented as mean with standard deviation from 3 technical replicates.

### Preclinical MPI of mouse organs

MPI of VivoTrax-labeled NK-92 cells administered to excised brains, legs, and lungs to mimic treatments for medulloblastoma, osteosarcoma, and lung metastases, are shown in **Figures 3-5**. For excised brains, no observable MPI signal was produced by control brains that received unlabeled cells (**Fig. 3a**). MPI signal is observed from the injection site for brains with 0.5 x 10^6^ or 1 x 10^6^ VivoTrax-labeled NK-92 cells, on the right side of the cerebellum (**Fig. 3b,c**). The integrated MPI signal from brains that received 1 x 10^6^ cells was ∼2.6x higher than 0.5 x 10^6^ cells, on average (n = 2) (**Fig. 3d**). 3D images show precise localization of 1x10^6^ NK-92 cells in the brain in axal, sagittal, and coronal views (**Fig. 3f-h**). The injection site localized by MPI was 1.6 mm right of midline (looking from hindbrain to forebrain), 1.2 mm caudal from lambda, and 3.1 mm down from cerebellar surface. Therefore, the injection site matched within 0.1-0.6 mm from the expected stereotactic injection site.

**Figure 3.**
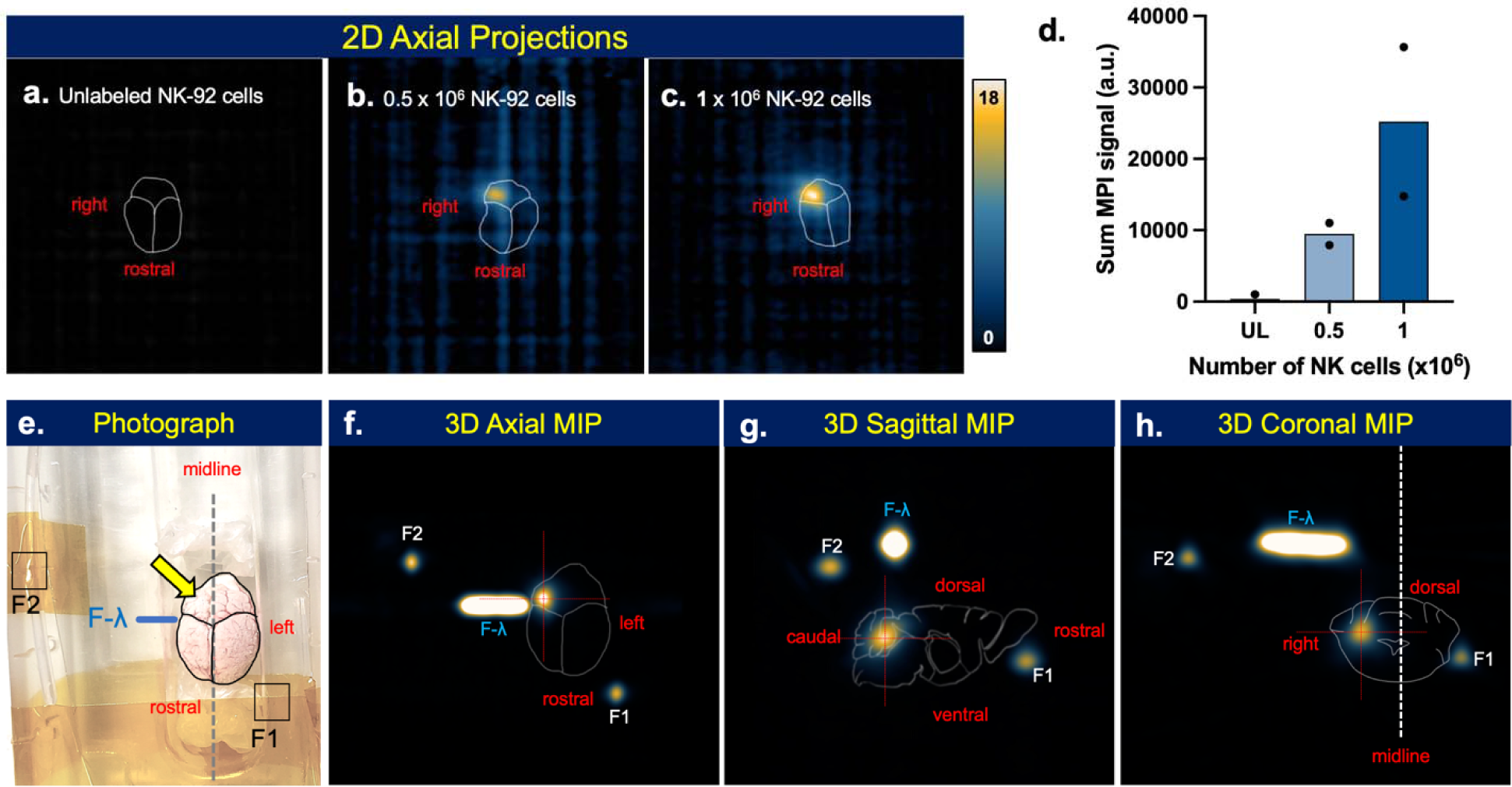
MPI detection, quantification, and localization of VivoTrax-labeled NK-92 cells implanted to mouse brain. A) No MPI signal was detected from brains injected with unlabeled cells (n = 4). MPI signal was detected on the right side of mouse brains with implanted VivoTrax-labeled NK-92 cells at a cell number of **b)** 0.5 x 10^6^ cells (n = 2) and **c)** 1x10^6^ cells (n = 2). Scale bars are 2 cm in length. **D)** Quantification of 2D images show increasing amount of MPI signal produced by unlabeled, 0.5x10^6^, and 1x10^6^ labeled NK-92 cells. Plots show mean with individual data points. **E**) Photograph of brain with fiducials for 3D imaging, with yellow arrow indicating the injection site. Three fiducials were included in the MPI field of view for localization; F1 placed at the left, rostral, ventral surface, F2 placed at the caudal, dorsal surface, and F- at lambda on the right side. 3D MPI of brain with 1x10^6^ VivoTrax-labeled NK-92 cells in the **f)** axial, **g)** sagittal, and **h)** coronal views. The injection site is indicated by red crosshairs. MIP = maximum intensity projection.

MPI signal from 2D projections is detected from the mid-tibia for legs with implanted VivoTrax-labeled NK-92 cells (**Fig. 4b,c**, n = 3), whereas control legs with unlabeled NK-92 cells show no observable signal above background noise (**Fig. 4a**, n = 6). Quantified MPI signal from 1x10^6^ NK-92 cells in tibia is 2.4x higher than 0.5x10^6^ cells, on average (** p < .001) (**Fig. 4d**). MPI signal from 0.5x10^6^ cells implanted in tibia is approximately 37x higher than background signals from unlabeled NK-92 cells (* p < .05). For 3D MPI, fiducials were aligned with the ankle and knee joints, and the medial surface of the leg for anatomical reference (**Fig. 4e**). Two focal MPI signals were identified in the tibia (**Fig. 4f,g**). The first signal was located at the target site; 8.6 mm caudal from knee joint (11.6 mm cranial from ankle joint), 2.4 mm lateral from the medial surface of the leg, and 5.0 mm dorsal of the ventral surface of the leg (marked in red). The second focal MPI signal was attributed to labeled cells that were implanted along the needle track, with maximum signal located at 5.2 mm caudal from knee joint (15.1 mm cranial from ankle joint), 1.9 mm lateral from the medial surface of the leg, and 2.5 mm dorsal of the ventral surface of the leg (marked in yellow). These focal regions are more prominently resolved in 3D compared to 2D (**Fig. 4h**).

**Figure 4.**
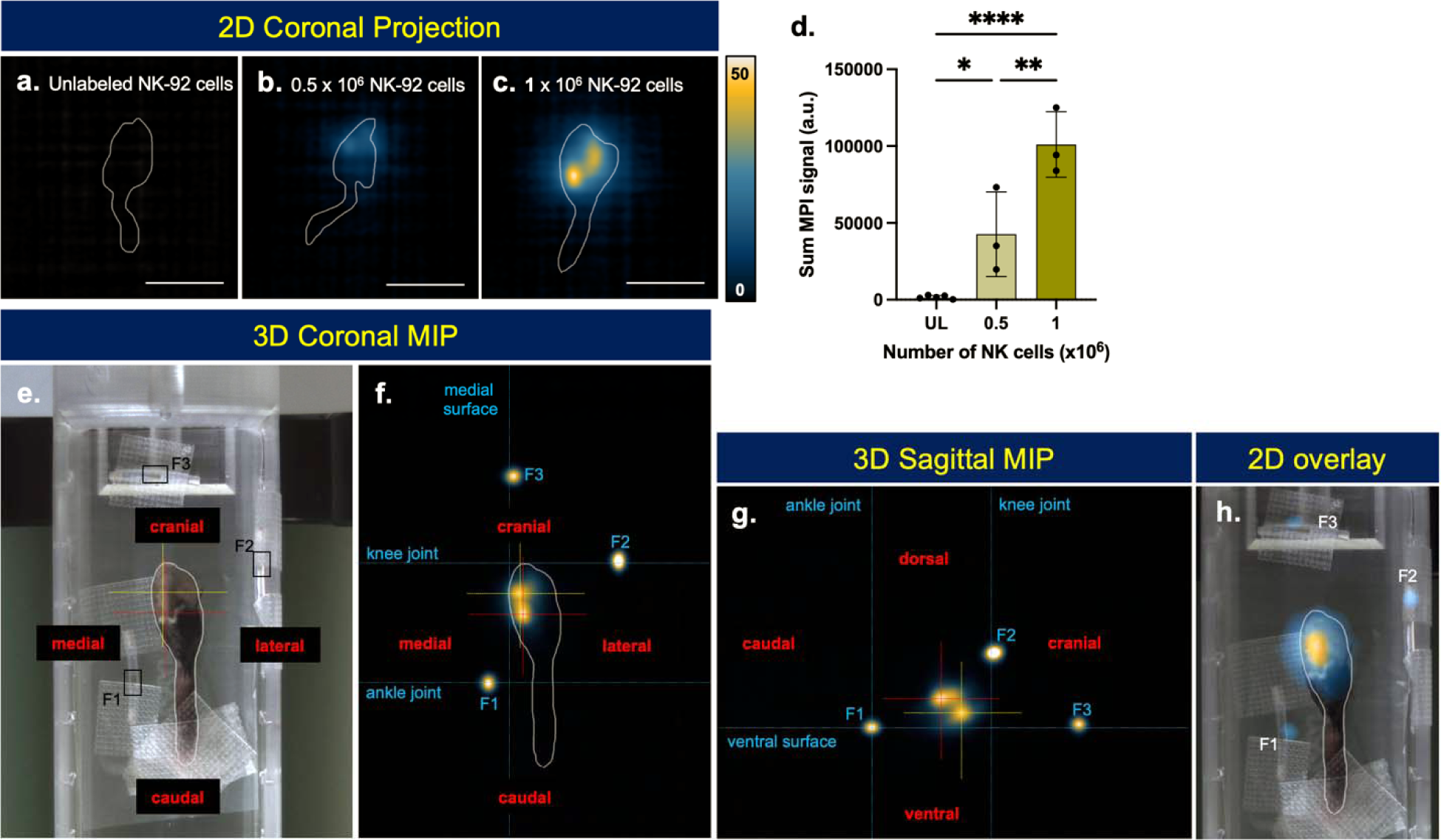
MPI detection, quantification, and localization of VivoTrax-labeled NK-92 cells implanted to mouse left tibias. A) No MPI signal was detected from tibias that received unlabeled NK-92 cells (n = 6). MPI signal was detected from tibias with implanted VivoTrax-labeled NK-92 cells at a cell number of **b)** 0.5 x 10^6^ cells (n = 3) and **c)** 1.0x10^6^ cells (n = 3). Scale bars are 2 cm in length. **D)** Quantification of 2D images shows significant differences in the amount of MPI signal from unlabeled, 0.5x10^6^ labeled and 1.0x10^6^ labeled NK-92 cells (* p < .05, ** p < .01, **** p < 0.0001). Plots show mean standard deviation with individual data points. **E)** Photograph of mouse tibia in MPI sample holder with 1x10^6^ implanted VivoTrax-labeled NK-92 cells. Three fiducials are placed for anatomical reference of the ankle joint (F1), knee joint (F2) and medial surface of leg (F3). Maximum intensity projections of 3D MPI in **f)** coronal and **g)** sagittal views identify 2 focal MPI signals in the tibia, marked by yellow and red crosshairs. **H)** Overlay of 2D MPI with photograph shows alignment of MPI signal with mouse leg.

**Figure 5.**
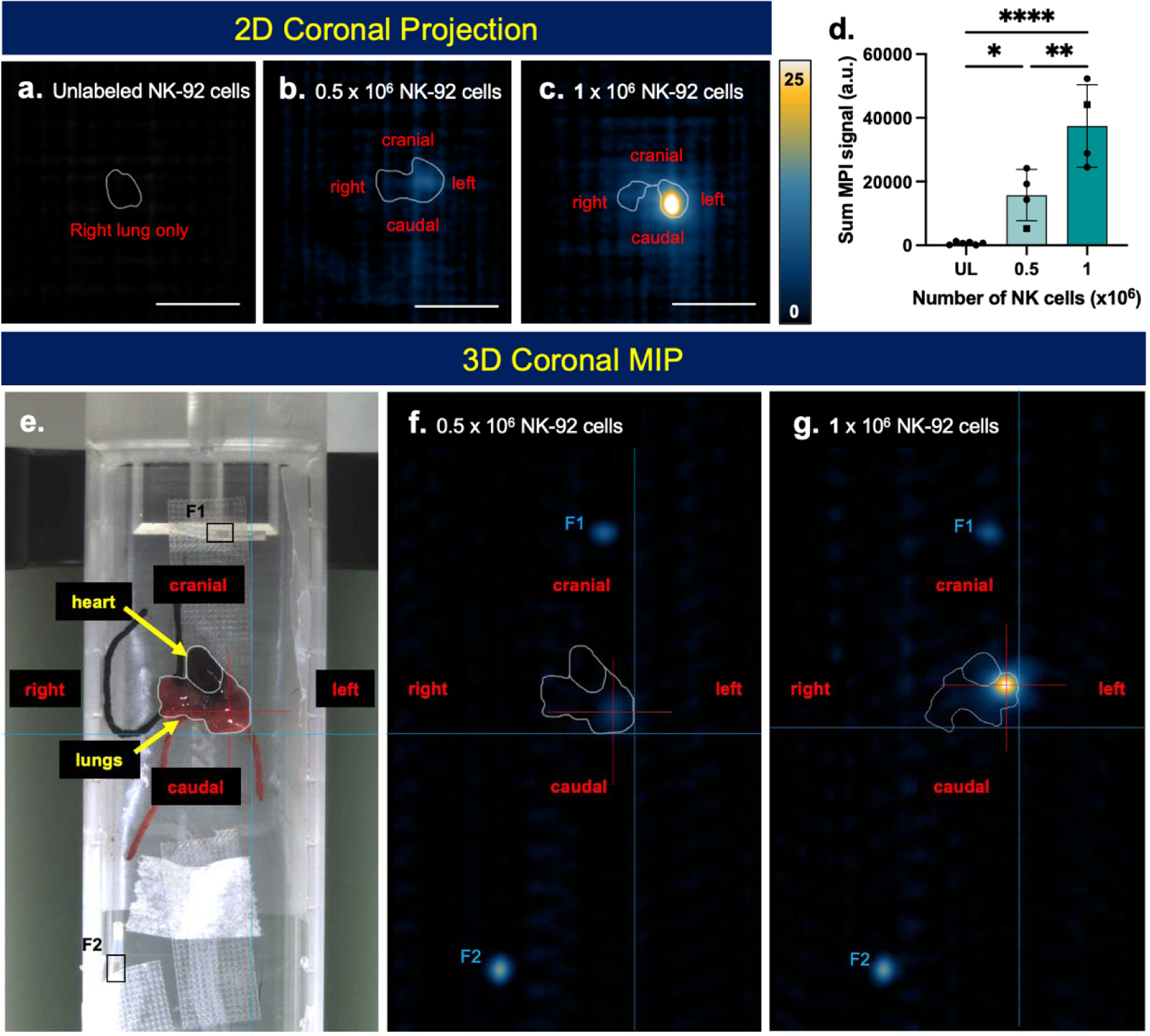
MPI detection, quantification, and localization of VivoTrax-labeled NK-92 cells implanted to mouse lungs. A) No MPI signal was detected from right lungs that received unlabeled NK-92 cells (n = 8). MPI signal was detected from left lungs with implanted VivoTrax-labeled NK-92 cells at a cell number of **b)** 0.5 x 10^6^ cells (n = 4) and **c)** 1.0x10^6^ cells (n = 4). Scale bars are 2 cm in length. Plots show mean standard deviation with individual data points. **D)** Quantification of 2D images show significant differences in the amount of MPI signal produced by unlabeled, 0.5x10^6^ labeled and 1.0x10^6^ labeled NK-92 cells (* p < .05, ** p < .01, **** p < 0.0001). **e)** Photograph of mouse lungs in MPI sample holder. Blue lines mark the left and caudal surface of the lungs. Coronal section from 3D MPI of **f)** 0.5 x 10^6^ NK-92 cells and **g)** 1.0 x 10^6^ NK-92 cells. Red targets mark the location of maximum MPI signal from NK-92 cell implantation.

MPI signal was present in the left lungs that received VivoTrax-labeled NK-92 cells and no MPI signal was present in the right lungs that received unlabeled NK-92 cells (**Fig. 5 a-c**). Quantified MPI signal from 1x10^6^ NK-92 cells in lung tissue is 2.8x higher than 0.5x10^6^ cells, on average (** p < .001) (**Fig. 5d**). MPI signal from 0.5x10^6^ cells implanted in lungs is approximately 22x higher than background signals from unlabeled NK-92 cells (* p < .05). Two sets of lungs, one with 0.5x10^6^ NK-92 cells (**Fig. 5f**) and the other with 1.0x10^6^ NK-92 cells (**Fig. 5g**), were imaged by 3D MPI surrounded by 3 fiducial markers placed on the base of the sample holder. Using the fiducials as position markers and photograph for anatomical reference, the location of MPI signal was determined for each lung. In **Fig 6f**, the MPI signal from labeled NK-92 cells in the left lung is 3.4 mm cranial from the caudal edge of the lung, 3.5 mm right of the left surface of the lung, and 1.0 mm deep from the anterior surface (base of sample holder). In **Fig 6g**, the MPI signal from labeled NK-92 cells in the left lung is 6.8 mm cranial from the caudal edge of the left lung, 2.0 mm right of the left surface of the lung, 1.3 mm deep from the anterior surface.

**Figure 6.**
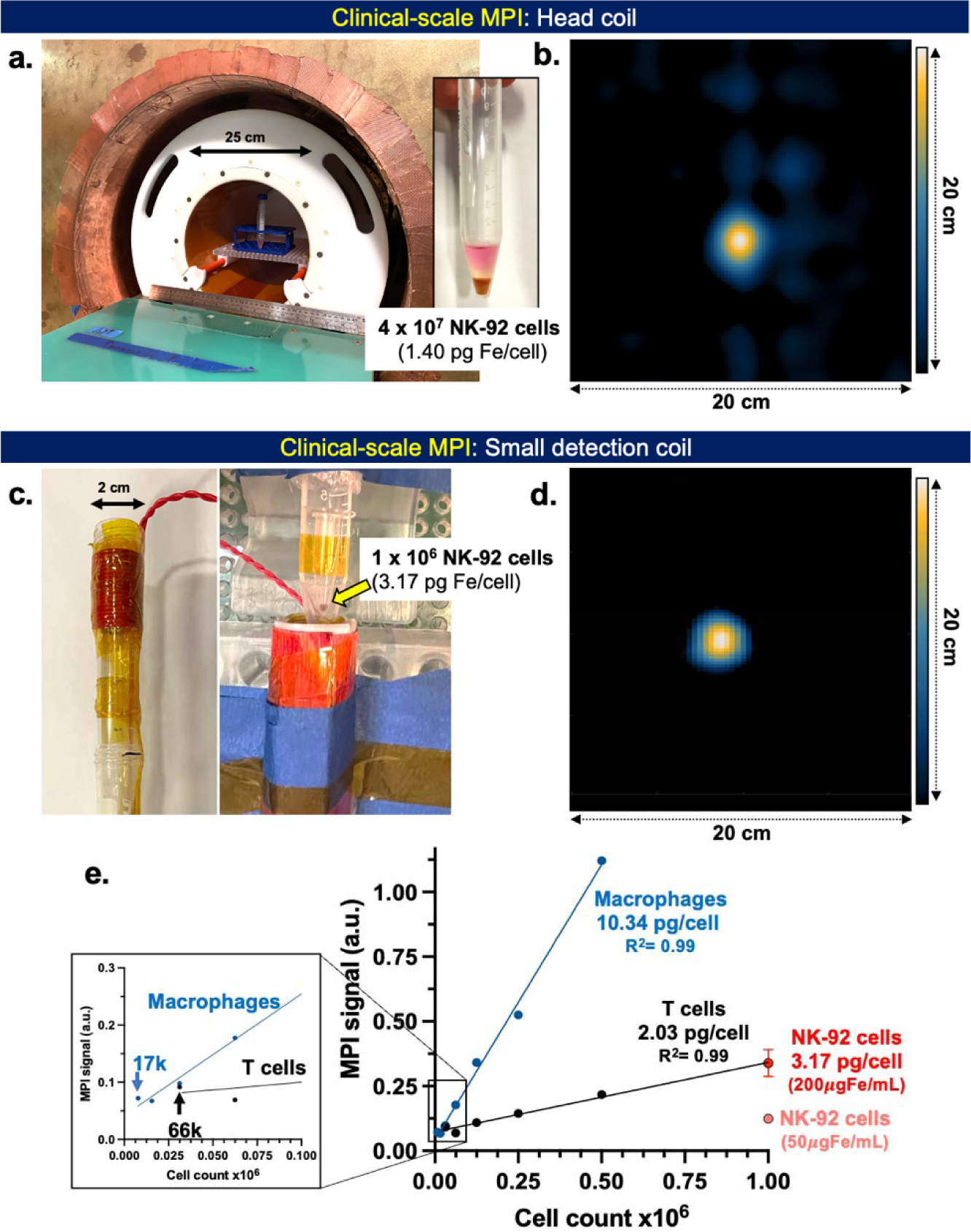
Clinical-scale MPI detection of VivoTrax-labeled NK-92 cells. (a) A VivoTrax-labeled NK-92 cell pellet with relevant clinical dose (4 x 10^7^ cells) was placed in the human MPI scanner (ID = 25 cm) and a 3D MPI image was acquired. (**b**) A 20 cm x 20 cm axial slice of the image is shown. **(c)** Cell pellets of labeled NK-92 cells, T cells, and macrophages ranging from 1.7 x 10^3^ cells – 1 x 10^6^ cells were imaged using a small detection coil with ID = 2 cm. **(d)** A 3D image was acquired of 1 x 10^6^ NK-92 cells, shown is a 20 cm x 20 cm axial slice. **(e)** The cellular limit of detection was established at 1.7 x 10^3^ macrophages or 6.6 x 10^3^ T cells. NK-92 cells (1 x 10^6^) produced similar signal as T cells of the same number.

### Clinical-scale MPI studies with NK-92 cells

We demonstrate detection of 40 million VivoTrax-labeled NK-92 cells using human MPI scanner (**Figure 6a**), producing an image with dimensions 20 x 20 x 3 cm and peak SNR = 110 (axial slice shown, **Figure 6b**). Using the small detection coil (**Figure 6c**), MPI of 1 million VivoTrax-labeled NK-92 cells (3.17 pg Fe/cell) was demonstrated (**Figure 6c,d**). Quantified MPI signal was similar for VivoTrax-labeled NK- 92 cells and T cells of the same number, as was seen on the preclinical MOMENTUM MPI scanner (**Figure 1d**). MPI signal from cell pellets was directly linear with number of VivoTrax-labeled T cells and macrophages (R^2^ = 0.99) (**Figure 6e**). The lowest number of cells detected on clinical-scale MPI using the small detection coil was 1.7x10^4^ macrophages (10.34 pg Fe/cell) and 6.6x10^4^ T cells (2.03 pg Fe/cell).

## Discussion

*In vivo* cell tracking has the potential to improve the accuracy and efficacy of cell therapies by allowing the monitoring of cell distribution and allowing the optimization of therapeutic cell therapies and protocols^46^. Indeed, our recently completed first Phase I clinical trial of pediatric patients with recurrent/refractory/metastatic fourth ventricular tumors, including medulloblastomas, highlights the need to address this clinical gap.^21^ Our trial’s primary endpoint demonstrated that intraventricular administration of autologous NK cells is feasible and safe. No adverse events were observed, with over 112 infusions of NK cells, up to doses of 10^9^ cells/m^2^. However, these treatments failed to demonstrate efficacy. The lack of an appropriate imaging biomarker prevents us from attributing failure to problems with NK cell delivery, localization to the tumor, or cytolytic activity. Additionally, our osteosarcoma pre-clinical studies demonstrated that a major determinant of NK cells therapeutic efficacy, when combined with IL-2, is the ability of NK cells to penetrate, proliferate and survive within the tumors.^47^ Others have similarly found that increased NK cell homing and infiltration within tumors is associated with improved tumor response and prognosis^48,49^, justifying the need for a non-invasive imaging biomarker able to identify and track ACT delivery. In a clinical setting, a new method of longitudinal cell tracking would for the first time provide clinicians with the ability to monitor ACT delivery to the region of interest in real time. It will also provide the ability to monitor tumor localization of ACT in the short-term. This new information would facilitate data-driven treatment decisions to inform clinical management.

The ideal cellular imaging technique provides sensitive and specific (unambiguous) detection of cells, with linear quantification and high resolution.^23,26,46^ Quantification is clinically valuable to determine the number of cells that were accurately delivered to a target location (or unintended location) and develop ACT dosing strategy. Direct cell labeling *in vitro* is a simple and common strategy employed for MRI cell tracking (using iron oxide nanoparticles and ^19^F-perfluorocarbons)^26^ and nuclear imaging (using ^111^In-oxine and ^89^Zr-oxine)^50^. ACTs have been safely tracked in humans using these cell labels^23,51–53^, however, no studies have been conducted to track NK cells in humans.

Iron-based MRI tracking has been the predominant approach for cell tracking in humans^23^. In MRI, iron oxide nanoparticles produce negative contrast and are sensitively detected due to what is known as the blooming artifact^54^. In one study, the distribution of iron-labeled stem cells was mapped for over 4 months following intraventricular injections^55^. However, negative signals for iron-based MRI cannot reliably be used to identify labeled cells due to other dark signals, such as anatomical features, air-filled lungs, or regions of hemorrhage^56^. Further, negative contrast is non-linear with iron quantity. Therefore, iron-based MRI is a sensitive cell tracking method but does not provide specific imaging signals to identify labeled cells or quantitation of cell number. ^19^F-perfluorocarbon-based MRI has been developed, which produces specific, hotspot images of ^19^F-labeled cells. Importantly, ^19^F MRI signal is directly linear with ^19^F concentration which can be used for quantification of cell number. Unfortunately, the drawback of this technique is low sensitivity, as noted by several preclinical studies with NK cells^27,28^ and a clinical study with dendritic cells^29^. ^19^F-MRI is also considered low resolution due to large voxel sizes needed to maximize sensitivity.

Nuclear imaging provides hotspot images with high sensitivity, however are limited by radiotracer half-life and it is challenging to quantify cell number.^46^ ^111^In-oxine (half-life = 2.8 days) has been used with SPECT imaging since the 1980s for leukocyte tracking,^53^ however, only offers low resolution^51^. ^89^Zr-oxine (half-life = 3.3 days) is gaining traction with PET imaging and has been used to track NK cells in macaques for 7 days.^57^ Unfortunately, nuclear imaging will not fulfill our clinical need to track delicate NK cells in pediatric populations and due to potential cytotoxic effects of ^89^Zr on cell activity even at low doses per cell.^30^

As we demonstrate in this work, MPI to addresses the limitations of current cellular imaging techniques. MPI specifically identifies the location of iron-labeled cells and produces a specific (hotspot) image^31–38^, similar to images obtained by ^19^F MRI and nuclear imaging. Here, we demonstrate that MPI also offers high resolution to determine cellular location. We show that MPI precisely localized labeled NK-92 cells within 1 mm of target injection sites, following technical injection to 3 different mouse organs (intracerebellar, intratibial, intrapulmonary). This was conducted using fiducial markers placed at anatomically significant locations but could be supplemented by co-registration with another anatomical image (e.g., CT or MRI). We have also demonstrated in this work that preclinical MPI sensitivity (∼ 3.1 x 10^4^ NK-92 cells at 3.14 µg Fe/10^6^ cells) is approaching clinical ^89^Zr-oxine PET (∼10^4^ cells at 15 kBq/10^6^ T cells).^50^

A major advantage for using cellular MPI is the ability to directly quantify iron from images, which can be related to the number of cells based on average cell labeling (pg/cell). We demonstrate strong linear relationships between MPI signal and number of NK-92 cells, T cells, and macrophages on both preclinical MPI and clinical-scale MPI. MPI signal was also linear with cell number for NK-92 cells in brain, tibia, and lung, irrespective of tissue depth and density. Therefore, we demonstrated the clinical requirement to accurately quantitate the number of NK-92 cells delivered to a target site (e.g., a tumor) using MPI. Further we demonstrate MPI has high specificity and resolution to determine if NK-92 cells were administered to the target location or otherwise identify a failed, or partially failed cell injection.

Our strategy for NK-92 cell labeling was co-incubation with VivoTrax in serum-free media^58–60^ and increasing VivoTrax concentration in the growth medium (50 vs. 200 µg Fe/mL). At the higher VivoTrax concentrations, there was both increased percentage of labeled cells and higher degree of labeling per cell. We did not observe any impairments to NK cell activity at these concentrations. Higher VivoTrax concentration for labeling was also associated with increased extracellular free iron (up to 40%). Based on previous studies, extracellular tracer is expected to quickly be cleared via the liver for ACT infusions^61^. Extracellular iron for ACTs administered to the cerebellum, tibia, or lungs may have different clearance kinetics of free tracer due to uptake and clearance by sentinel phagocytic cells. This will be explored in future studies. We achieved similar cell uptake, MPI signal, and limit of detection for NK-92 cells (3.17 pg Fe/cell) and Jurkat T cells (2.03 pg/cell) which reflects their similar endocytic capacity.

Direct cell labeling approaches can only reliably enable short-term tracking and quantification of cells. It is currently unknown how long NK cells could be tracked with MPI due to cell proliferation and dilution of iron oxide nanoparticles amongst cell progeny. For long-term cell tracking, other imaging approaches using reporter genes are being explored with MRI and PET^62–64^, however these involve genetic manipulations of ACTs. A recent review compares indirect and direct cell labeling.^65,66^

Our future work will be focused on labeling primary allogenic NK cells expanded from patient donors for MPI. In our previous Phase I study of intraventricular infusions of autologous ex vivo expanded NK cells, the maximum tolerated dose we studied in children with recurrent medulloblastoma and ependymoma was 300 x 10^6^ cells.^21^ This dose was deemed safe and will be the starting dose of our upcoming trials. In this paper we successfully imaged 40 x 10^6^ VivoTrax-labeled NK cells on a clinical-scale MPI, representing 13.3% of an infused dose. While VivoTrax is not FDA-approved for human use, other carboxydextran-coated superparamagnetic iron oxide nanoparticles are FDA-approved and could be used off-label for cell labeling and MPI tracking. As iron-based MRI has been used in 8 cell tracking studies in humans^23^, we anticipate that iron-based cell tracking with MPI to also be safe. As a clinical imaging biomarker, MPI could play a role in clinical trials, inform clinical decision-making, and improve patient outcomes.

## Conclusion

Allogenic NK cell therapies are promising for treatment of medulloblastoma and osteosarcoma, but cell tracking is needed to provide data-driven development of new cell therapies and, ultimately, treatment decisions. Here we demonstrated protocol to label NK-92 cells with iron oxide nanoparticles, resulting in detection using both preclinical MPI and clinical-scale MPI. Labeled NK-92 cells were precisely localized and quantified by preclinical MPI following administration to the cerebellum, tibia, and lungs. Importantly, NK-92 cells maintained their viability and cytolysis activity against medulloblastoma and osteosarcoma cell lines with VivoTrax labeling. MPI can provide sensitive and quantitative information on NK-92 cell delivery, therefore has the potential to resolve a significant unmet clinical need to track NK cell treatments in patients.

## Acknowledgements

We gratefully acknowledge support from Addis Faith Foundation (V.G.) and CRPIT: RP200223 (K.S., V.G.). Authors O.C.S., K.G., B.F., A.R.M., E.E.M., T.S., P.K., D.T., P.W.G., and J.M.G. report relationship with Magnetic Insight Inc. that includes: employment and stock ownership. These authors have contributed to study design, writing of the report, and in the decision to submit the paper for publication.

## Notes

### Competing Interest Statement

Olivia C. Sehl, Kelvin Guo, Benjamin Fellows, A. Rahman Mohtasebzadeh, Erica E. Mason, Toby Sanders, Petrina Kim, David Trease, Patrick W. Goodwill, and Joan M Greve report relationship with Magnetic Insight Inc. that includes: employment and stock ownership.

## References

1. Suk Y, Gwynne WD, Burns I, Venugopal C, Singh SK. Childhood Medulloblastoma: An Overview. 2022. pp. 1–12.

2. Mirabello, L., Troisi, R.J. and Savage, S.A., 2009. Osteosarcoma incidence and survival rates from 1973 to 2004: data from the Surveillance, Epidemiology, and End Results Program. Cancer: Interdisciplinary International Journal of the American Cancer Society, 115(7), pp.1531–1543.

3. Moreno, F., Cacciavillano, W., Cipolla, M., Coirini, M., Streitenberger, P., López Martí, J., Palladino, M., Morici, M., Onoratelli, M., Drago, G. and Schifino, A., 2017. Childhood osteosarcoma: Incidence and survival in Argentina. Report from the national pediatric cancer registry, ROHA network 2000–2013. Pediatric blood & cancer, 64(10), p.e26533.

4. Gok Durnali, A., Paksoy Turkoz, F., Ardic Yukruk, F., Tokluoglu, S., Yazici, O.K., Demirci, A., Bal, O., Gundogdu Buyukbas, S., Esbah, O., Oksuzoglu, B. and Alkis, N., 2016. Outcomes of adolescent and adult patients with lung metastatic osteosarcoma and comparison of synchronous and metachronous lung metastatic groups. PLoS One, 11(5), p.e0152621.

5. Epperly, R., Gottschalk, S. and Velasquez, M.P., 2020. Harnessing T cells to target pediatric acute myeloid leukemia: CARs, BiTEs, and beyond. Children, 7(2), p.14

6. Chang, T.C., Carter, R.A., Li, Y., Li, Y., Wang, H., Edmonson, M.N., Chen, X., Arnold, P., Geiger, T.L., Wu, G. and Peng, J., 2017. The neoepitope landscape in pediatric cancers. Genome medicine, 9, pp.1–12.

7. Thomas, B.C., Staudt, D.E., Douglas, A.M., Monje, M., Vitanza, N.A. and Dun, M.D., 2023. CAR T cell therapies for diffuse midline glioma. Trends in cancer.

7. DeRenzo, C., Krenciute, G. and Gottschalk, S., 2018. The landscape of CAR T cells beyond acute lymphoblastic leukemia for pediatric solid tumors. American Society of Clinical Oncology Educational Book, 38, pp.830–837.

8. Zhang, J. and Wang, T., 2021. Immune cell landscape and immunotherapy of medulloblastoma. Pediatric Investigation, 5(04), pp.299–309.

9. Shimasaki, N., Jain, A. and Campana, D., 2020. NK cells for cancer immunotherapy. Nature reviews Drug discovery, 19(3), pp.200–218.

10. Liu S, Galat V, Galat4 Y, Lee YKA, Wainwright D, Wu J. NK cell-based cancer immunotherapy: from basic biology to clinical development. J Hematol Oncol. 2021;14: 7.

11. Myers JA, Miller JS. Exploring the NK cell platform for cancer immunotherapy. Nat Rev Clin Oncol. 2021;18(2):85–100.

12. Toffoli EC, Sheikhi A, Höppner YD, de Kok P, Yazdanpanah-Samani M, Spanholtz J, Verheul HMW, van der Vliet HJ, de Gruijl TD. Natural Killer Cells and Anti-Cancer Therapies: Reciprocal Effects on Immune Function and Therapeutic Response. Cancers (Basel). 2021;13: 711.

13. Cheng M, Chen Y, Xiao W, Sun R, Tian Z. NK cell-based immunotherapy for malignant diseases. Cell Mol Immunol. 2013;10: 230–252.

14. Shimasaki N, Jain A, Campana D. NK cells for cancer immunotherapy. Nat Rev Drug Discov. 2020;19: 200–218.

15. Valipour B, Velaei K, Abedelahi A, Karimipour M, Darabi M, Charoudeh HN. NK cells: An attractive candidate for cancer therapy. J Cell Physiol. 2019;234(11):19352–65.

16. Tarek, N. and Lee, D.A., 2014. Natural killer cells for osteosarcoma. Current Advances in Osteosarcoma, pp.341–353.

17. Buddingh, E.P., Schilham, M.W., Ruslan, S.E.N., Berghuis, D., Szuhai, K., Suurmond, J., Taminiau, A.H., Gelderblom, H., Egeler, R.M., Serra, M. and Hogendoorn, P.C., 2011. Chemotherapy-resistant osteosarcoma is highly susceptible to IL-15-activated allogeneic and autologous NK cells. Cancer immunology, immunotherapy, 60, pp.575–586.

19. Laskowski, T.J., Biederstädt, A. and Rezvani, K., 2022. Natural killer cells in antitumour adoptive cell immunotherapy. Nature Reviews Cancer, 22(10), pp.557–575.

18. Denman CJ, Senyukov V V., Somanchi SS, Phatarpekar P V., Kopp LM, Johnson JL, Singh H, Hurton L, Maiti SN, Huls MH, Champlin RE, Cooper LJN, Lee DA. Membrane-Bound IL-21 Promotes Sustained Ex Vivo Proliferation of Human Natural Killer Cells. PLoS One. 2012;7: e30264.

19. Khatua S, Cooper LJN, Sandberg DI, Ketonen L, Johnson JM, Rytting ME, Liu DD, Meador H, Trikha P, Nakkula RJ, Behbehani GK, Ragoonanan D, Gupta S, Kotrotsou A, Idris T, Shpall EJ, Rezvani K, Colen R, Zaky W, Lee DA, Gopalakrishnan V. Phase I study of intraventricular infusions of autologous ex vivo expanded NK cells in children with recurrent medulloblastoma and ependymoma. Neuro Oncol. 2020;22: 1214–1225.

20. Chawla, S., Shehu, V., Gupta, P.K., Nath, K. and Poptani, H., 2021. Physiological imaging methods for evaluating response to immunotherapies in glioblastomas. International Journal of Molecular Sciences, 22(8), p.3867

21. Bulte, J. W., & Daldrup-Link, H. E. (2018). Clinical tracking of cell transfer and cell transplantation: trials and tribulations. Radiology, 289(3), 604–615.

22. Malosio ML, Esposito A, Brigatti C, et al. (2015). MR imaging monitoring of iron-labeled pancreatic islets in a small series of patients: islet fate in successful, unsuccessful, and autotransplantation. Cell Transplant 24(11):2285–2296.

23. Ahrens ET, Flores R, Xu H, Morel PA. In vivo imaging platform for tracking immunotherapeutic cells. Nat Biotechnol. 2005;23: 983–987.

24. Ahrens ET, Bulte JWM. Tracking immune cells in vivo using magnetic resonance imaging. Nature Reviews Immunology 2013 13:10. 2013;13: 755–763.

25. Kennis, B.A., Michel, K.A., Brugmann, W.B., Laureano, A., Tao, R.H., Somanchi, S.S., Einstein, S.A., Bravo-Alegria, J.B., Maegawa, S., Wahba, A. and Kiany, S., 2019. Monitoring of intracerebellarly-administered natural killer cells with fluorine-19 MRI. Journal of neuro-oncology, 142, pp.395–407.

28. Bouchlaka, Myriam N., Kai D. Ludwig, Jeremy W. Gordon, Matthew P. Kutz, Bryan P. Bednarz, Sean B. Fain, and Christian M. Capitini. “19F-MRI for monitoring human NK cells in vivo.” Oncoimmunology 5, no. 5 (2016): e1143996.

26. Ahrens, Eric T., Brooke M. Helfer, Charles F. O’Hanlon, and Claudiu Schirda. “Clinical cell therapy imaging using a perfluorocarbon tracer and fluorine-19 MRI.” Magnetic resonance in medicine 72, no. 6 (2014): 1696–1701.

27. Jacob, Jacinta, Alessia Volpe, Qi Peng, Robert I. Lechler, Lesley A. Smyth, Giovanna Lombardi, and Gilbert O. Fruhwirth. “Radiolabelling of Polyclonally Expanded Human Regulatory T Cells (Treg) with 89Zr-oxine for Medium-Term In Vivo Cell Tracking.” Molecules 28, no. 3 (2023): 1482.

28. Zheng, et al. Quantitative MPI monitors the transplantation, biodistribution, and clearance of stem cells in vivo. Theranostics, 2016. 6(3):291–301.

29. Wang, Q., Ma, X., Liao, H., Liang, Z., Li, F., Tian, J., & Ling, D. (2020). Artificially Engineered Cubic Iron Oxide Nanoparticle as a High-Performance Magnetic Particle Imaging Tracer for Stem Cell Tracking. ACS nano, 14(2), 2053–2062.

30. Fink, Corby, Julia J. Gevaert, John W. Barrett, Jimmy D. Dikeakos, Paula J. Foster, and Gregory A. Dekaban. “In vivo tracking of adenoviral-transduced iron oxide-labeled bone marrow-derived dendritic cells using magnetic particle imaging.” Eur radio express (2023).

31. Bulte, Jeff WM, and Ali Shakeri-Zadeh. “In vivo MRI tracking of tumor vaccination and antigen presentation by dendritic cells.” Molecular Imaging and Biology (2022): 1–10.

32. Makela, A. V., Schott, M. A., Sehl, O. C., Gevaert, J. J., Foster, P. J., & Contag, C. H. (2022). Tracking the fates of iron-labeled tumor cells in vivo using Magnetic Particle Imaging. Nanoscale Advances, 4(17), 3617–3623.

33. Wu, Lyndia C., Yanrong Zhang, Gary Steinberg, H. Qu, S. Huang, M. Cheng, T. Bliss et al. “A review of magnetic particle imaging and perspectives on neuroimaging.” American Journal of Neuroradiology 40, no. 2 (2019): 206–212.

34. Rivera-Rodriguez, Angelie, Lan B. Hoang-Minh, Andreina Chiu-Lam, Nicole Sarna, Leyda Marrero-Morales, Duane A. Mitchell, and Carlos M. Rinaldi-Ramos. “Tracking adoptive T cell immunotherapy using magnetic particle imaging.” Nanotheranostics 5, no. 4 (2021): 431.

35. Wu, Wei Emma, Edwin Chang, Linchun Jin, Shiqin Liu, Ching-Hsin Huang, Rozy Kamal, Tie Liang et al. “Multimodal In Vivo Tracking of Chimeric Antigen Receptor T Cells in Preclinical Glioblastoma Models.” Investigative Radiology 58, no. 6 (2023): 388–395.

36. Mattingly, E., Mason, E., Sliwiak, M. and Wald, L.L., 2022. Drive and receive coil design for a human-scale MPI system. International Journal on Magnetic Particle Imaging IJMPI, *8*(1 Suppl 1).

37. Gräser, M., Thieben, F., Szwargulski, P., Werner, F., Gdaniec, N., Boberg, M., Griese, F., Möddel, M., Ludewig, P., Van De Ven, D. and Weber, O.M., 2019. Human-sized magnetic particle imaging for brain applications. Nature communications, 10(1), p.1936.

38. Vogel, P., Rückert, M., Greiner, C., Günther, J., Reichl, T., Kampf, T., Bley, T., Behr, V. and Herz, S., 2022. iMPI–portable human-sized Magnetic Particle Imaging Scanner for real-time endovascular Interventions.

39. Le, T.A., Bui, M.P., Gadelmowla, K.M., Oh, S. and Yoon, J., 2023. First Human-scale Magnetic Particle Imaging System with Superconductor. International Journal on Magnetic Particle Imaging IJMPI, *9*(1 Suppl 1).

40. Mason EE, Barcikowski E, Carl J, Chandeeing M, Davidson B, Fellows B, Fetsch W, Fields K, Greve J, Konkle JJ, Mattingly E, Sanders T, Sehl OC, Trease D, Truxal A, Weyhmiller M, Goodwill P. Preliminary Results: Large Bore Clinical MPI System Imaging Human Head-sized FOVs. International Journal on Magnetic Particle Imaging (2024).

44. 41. International Organization for Standardization. (2018). Nanotechnologies – In vitro MTS assay for measuring the cytotoxic effect of nanoparticles (ISO Standard No. 19007:2018).

42. Sehl, O.C., Tiret, B., Berih, M.A., Makela, A.V., Goodwill, P.W. and Foster, P.J., 2022. MPI region of interest (ROI) analysis and quantification of iron in different volumes. International Journal on Magnetic Particle Imaging, 8(1).

46. Helfer, B.M., Ponomarev, V., Patrick, P.S., Blower, P.J., Feitel, A., Fruhwirth, G.O., Jackman, S., Mouries, L.P., Park, M.V., Srinivas, M. and Stuckey, D.J., 2021. Options for imaging cellular therapeutics in vivo: a multi-stakeholder perspective. Cytotherapy, 23(9)

43. Guma, S.R., Lee, D.A., Ling, Y., Gordon, N. and Kleinerman, E.S., 2014. Aerosol interleukin-2 induces natural killer cell proliferation in the lung and combination therapy improves the survival of mice with osteosarcoma lung metastasis. Pediatric blood & cancer, 61(8), pp.1362–1368.

44. Nersesian, S., Schwartz, S.L., Grantham, S.R., MacLean, L.K., Lee, S.N., Pugh-Toole, M. and Boudreau, J.E., 2021. NK cell infiltration is associated with improved overall survival in solid cancers: A systematic review and meta-analysis. Translational oncology, 14(1), p.100930.

45. Nayyar, G., Chu, Y. and Cairo, M.S., 2019. Overcoming resistance to natural killer cell based immunotherapies for solid tumors. Frontiers in oncology, 9, p.51.

46. Lechermann, Laura M., Doreen Lau, Bala Attili, Luigi Aloj, and Ferdia A. Gallagher. “In vivo cell tracking using PET: Opportunities and challenges for clinical translation in oncology.” Cancers 13, no. 16 (2021): 4042.

47. De Vries, I.J.M., Lesterhuis, W.J., Barentsz, J.O., Verdijk, P., Van Krieken, J.H., Boerman, O.C., Oyen, W.J., Bonenkamp, J.J., Boezeman, J.B., Adema, G.J. and Bulte, J.W., 2005. Magnetic resonance tracking of dendritic cells in melanoma patients for monitoring of cellular therapy. Nature biotechnology, 23(11), pp.1407–1413.

48. Lapi, S., McConathy, J., Jeffers, C., Bartels, J., Houson, H., White, S. and Younger, J., 2022. First-in-Human Imaging of 89Zr-oxine Labelled Autologous Leukocytes in Healthy Volunteers.

49. McAfee, J.G.; Samin, A. In-111 labeled leukocytes: A review of problems in image interpretation. Radiology 1985, 155, 221–229.

50. Shapiro, E.M., Sharer, K., Skrtic, S. and Koretsky, A.P., 2006. In vivo detection of single cells by MRI. Magnetic Resonance in Medicine: An Official Journal of the International Society for Magnetic Resonance in Medicine, 55(2), pp.242–249.

55. Janowski, M., Walczak, P., Kropiwnicki, T., Jurkiewicz, E., Domanska-Janik, K., Bulte, J.W., Lukomska, B. and Roszkowski, M., 2014. Long-term MRI cell tracking after intraventricular delivery in a patient with global cerebral ischemia and prospects for magnetic navigation of stem cells within the CSF. PloS one, 9(6), p.e97631.

51. Iv, M., Samghabadi, P., Holdsworth, S., Gentles, A., Rezaii, P., Harsh, G., Li, G., Thomas, R., Moseley, M., Daldrup-Link, H.E. and Vogel, H., 2019. Quantification of macrophages in high-grade gliomas by using ferumoxytol-enhanced MRI: a pilot study. Radiology, 290(1), pp.198–206.

52. Sato, N., Stringaris, K., Davidson-Moncada, J.K., Reger, R., Adler, S.S., Dunbar, C., Choyke, P.L. and Childs, R.W., 2020. In vivo tracking of adoptively transferred natural killer cells in rhesus macaques using 89Zirconium-oxine cell labeling and PET imaging. Clinical Cancer Research, 26(11), pp.2573–2581.

53. Santos, António JM, and Emmanuel Boucrot. “Probing endocytosis during the cell cycle with minimal experimental perturbation.” Clathrin-Mediated Endocytosis: Methods and Protocols (2018): 23–35.

54. Boddington, Sophie, Tobias D. Henning, Elizabeth J. Sutton, and Heike E. Daldrup-Link. “Labeling stem cells with fluorescent dyes for non-invasive detection with optical imaging.” JoVE (Journal of Visualized Experiments) 14 (2008): e686.

55. Zhu, Ying, Wenxin Li, Qingnuan Li, Yuguo Li, Yufeng Li, Xiaoyong Zhang, and Qing Huang. “Effects of serum proteins on intracellular uptake and cytotoxicity of carbon nanoparticles.” Carbon 47, no. 5 (2009): 1351–135

56. Liu, S., Chiu-Lam, A., Rivera-Rodriguez, A., DeGroff, R., Savliwala, S., Sarna, N. and Rinaldi-Ramos, C.M., 2021. Long circulating tracer tailored for magnetic particle imaging. Nanotheranostics, 5(3), p.348.

57. Nyström, N.N., McRae, S.W., Martinez, F.M., Kelly, J.J., Scholl, T.J. and Ronald, J.A., 2023. A Genetically Encoded Magnetic Resonance Imaging Reporter Enables Sensitive Detection and Tracking of Spontaneous Metastases in Deep Tissues. Cancer Research, 83(5), pp.673–685

58. Jacob, J., Nadkarni, S., Volpe, A., Peng, Q., Tung, S.L., Hannen, R.F., Mohseni, Y.R., Scotta, C., Marelli-Berg, F.M., Lechler, R.I. and Smyth, L.A., 2021. Spatiotemporal in vivo tracking of polyclonal human regulatory T cells (Tregs) reveals a role for innate immune cells in Treg transplant recruitment. Molecular Therapy-Methods & Clinical Development, 20, pp.324–336.

59. Keu K.V., Witney T., Yaghoubi S., Rosenberg J., Kurien A., Magnusson R., Williams J., Habte F., Wagner J.R., Forman S., et al. Reporter gene imaging of targeted T cell immunotherapy in recurrent glioma. Sci. Transl. Med. 2017;9:eaag2196.

60. Bulte, J.W., 2024. Direct versus Indirect Labeling for Chimeric Antigen Receptor T-Cell Tracking Using PET. Radiology, 310(2), p.e240241.

61. Galli, Filippo, Michela Varani, Chiara Lauri, Guido Gentiloni Silveri, Livia Onofrio, and Alberto Signore. “Immune cell labelling and tracking: implications for adoptive cell transfer therapies.” EJNMMI Radiopharmacy and Chemistry 6, no. 1 (2021): 1–19.

