## Supplementary Figure S1 for "Labeling Natural Killer cells with superparamagnetic iron oxide nanoparticles for detection by preclinical and clinical-scale magnetic particle imaging"

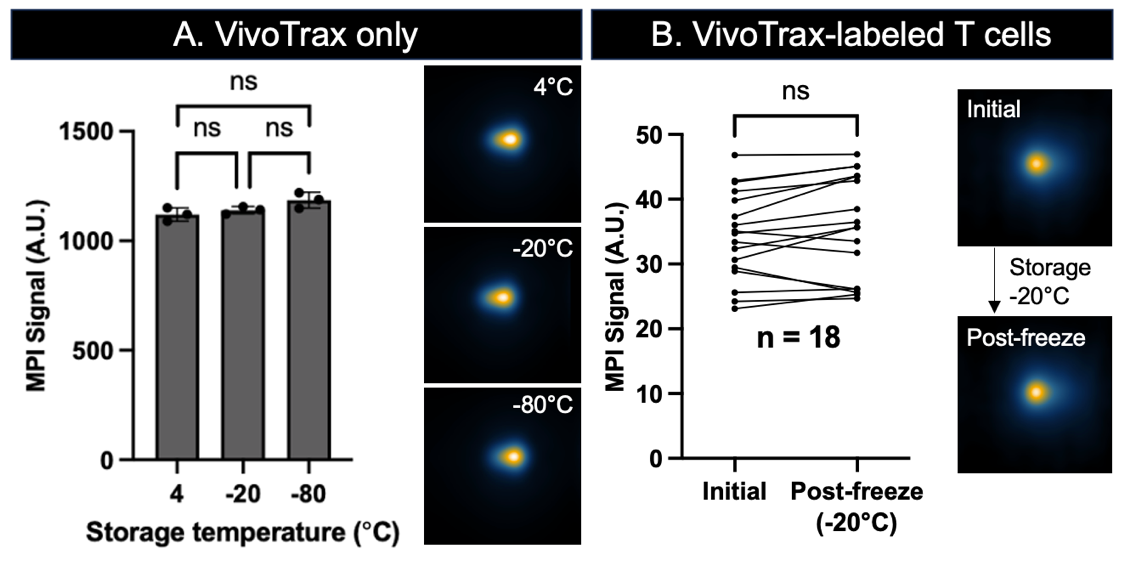

**Supplementary Figure S1. MPI signal was maintained in T cells that experienced a freeze-thaw cycle. A.** MPI signal was not significantly different for 5 µg VivoTrax stored at 4, -20, or -80 $^{\circ}$C for 24 hours (n = 3, ANOVA, p > .05). **B.** MPI signal was maintained in T cells that experienced a freeze-thaw cycle. MPI images show VivoTrax-labeled T cells before and after -20 $^{\circ}$C storage (n = 18, paired t-test, p > .05). This result provides valuable flexibility in the workflow by allowing for labeling at a central location and shipping of labeled, frozen cells to the point of use.
